# A functional approach to homeostatic regulation

**DOI:** 10.1101/2024.05.22.595277

**Authors:** Clemente F. Arias, Francisco J. Acosta, Federica Bertocchini, Cristina Fernández-Arias

**Affiliations:** Grupo Interdisciplinar de Sistemas Complejos de Madrid (GISC), 28040 Madrid, Spain; Departamento de Ecología, Universidad Complutense de Madrid, 28040 Madrid, Spain; Plasticentropy, 51100 Reims, France; Departamento de Inmunología, Facultad de Medicina, Universidad Complutense de Madrid, 28040 Madrid, Spain

**Keywords:** Homeostasis, physiology, metabolism, aconitase, hepcidin, intracellular iron homeostasis, systemic iron homeostasis, insulin, blood glucose homeostasis, Hypoxia-inducible factors (HIFs), intracellular oxygen homeostasis, control theory

## Abstract

In this work, we present a novel modeling framework for understanding the dynamics of homeostatic regulation. Inspired by engineering control theory, this framework incorporates unique features of biological systems. First, biological variables often play physiological roles, and taking this functional context into consideration is essential to fully understand the goals and constraints of homeostatic regulation. Second, biological signals are not abstract variables, but rather material molecules that may undergo complex turnover processes of synthesis and degradation. We suggest that the particular nature of biological signals may condition the type of information they can convey, and their potential role in shaping the dynamics and the ultimate purpose of homeostatic systems. We show that the dynamic interplay between regulated variables and control signals is a key determinant of biological homeostasis, challenging the necessity and the convenience of strictly extrapolating concepts from engineering control theory in modeling the dynamics of homeostatic systems. This work provides an alternative, unified framework for studying biological regulation and identifies general principles that transcend molecular details of particular homeostatic mechanisms. We show how this approach can be naturally applied to apparently different regulatory systems, contributing to a deeper understanding of homeostasis as a fundamental process in living systems.

## Introduction

Homeostasis denotes the ability of living systems to maintain internal stability under the influence of external factors [1]. The origin of this concept lies in the observation that key physiological variables, such as body temperature, blood pH, or blood glucose levels, remain within narrow ranges in a vast array of circumstances [1, 2]. However, in recent decades, the meaning of ‘homeostasis’ has broadened to encompass the physiological processes that allow biological systems to cope with unexpected changes in either internal or external conditions [3–5].

Although the notion of homeostasis pervades biology, it lacks a rigorous formalism. This has motivated the use of terminology from engineering control theory to describe homeostatic processes [6–12]. From this approach, the regulation of a given target variable relies on three primary elements: sensors, controllers, and actuators. Sensors detect the current value of the target variable and compare it with a reference value, normally labeled as ‘set point’ [13]. Controllers process the information received from sensors and generate a signal that induces the actuators to adjust the system’s behavior, thereby bringing the value of the regulated variable closer to the set point (Fig. 1.A). Describing homeostatic regulation in these terms is straightforward [14] (see Figs. 1.B,C).

**Figure 1.**
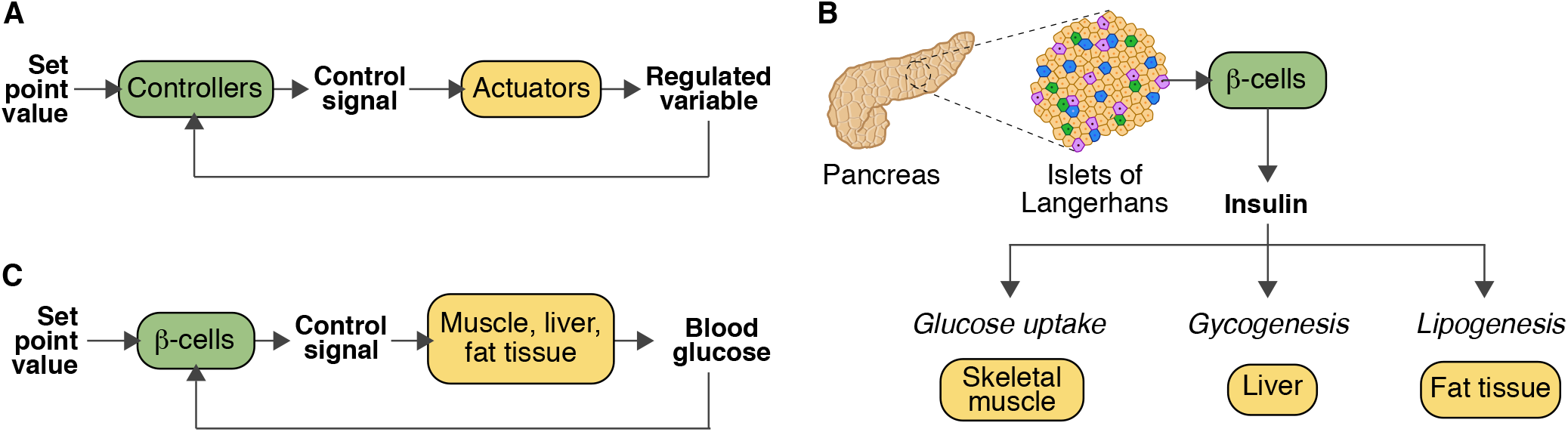
Application of control theory to biological regulation. A) Block diagram of a typical engineering control system. B) Regulation of blood glucose by insulin. Pancreatic *β* -cells continuously monitor blood glucose levels and react to hyperglycemia by secreting insulin into the bloodstream. Insulin targets a variety of tissues across the body, including muscle, liver, and adipose tissue, promoting the uptake, storage, and utilization of glucose, thus reducing the concentration of glucose in the blood. C) In terms of control theory, pancreatic *β* -cells act simultaneously as sensors, detecting whether glucose levels exceed a critical threshold (the set point), and controllers, using insulin to orchestrate a coordinated systemic response aimed at lowering blood glucose through its action on a variety of actuators scattered throughout the organism. This mode of action defines a feedback mechanism that brings the system toward its predefined set point.

In control theory, the set point defines the desired state for the system, and the deviation from this state is termed the system’s error [13]. Accordingly, changes in the value of the regulated variable are considered as unwanted disturbance signals and sources of error that interfere with the function of the system [13]. The goal of regulation is to minimize or eliminate these disturbances, ensuring that the system remains as close as possible to its preferred state, given by set point. This view of regulation can be easily extrapolated to biological homeostasis. The concept of set point neatly captures the tendency of some biological variables to exhibit a relatively constant value [7], which is widely assumed as the hallmark of homeostasis [1]. Moreover, the view of changes in regulated variables as perturbations that disrupt the optimal state of the system was already explicit in Bernard’s and Cannon’s seminal works [15], and it is currently widespread in homeostasis studies [16–19].

In our opinion, despite the compelling analogies between control systems in engineering and biology, understanding homeostatic regulation as a response to perturbations that deviate regulated variables from their set points poses some major issues. Whereas this interpretation of homeostasis may be useful for traits such as blood pH or body temperature, it may not be appropriate for many other physiological variables, such as blood glucose or oxygen levels. From a functional viewpoint, assuming that the main role of blood glucose is to achieve a fixed steady state value is misleading. The role of glucose is better understood in relation to the metabolic demands of the body tissues. Glucose consumption and uptake are integral parts of this metabolic function, even if they may cause marked fluctuations that move blood glucose levels away from the set point. Such fluctuations in the value of a regulated variable should not be viewed as detrimental perturbations of the system’s optimal steady state; rather, they should be understood as normal aspects of homeostatic dynamics.

In this work, we formulate a conceptual framework to study homeostasis. Central in our approach is the assumption that regulated variables normally have physiological functions [20]. We suggest that taking this broader functional background into account is essential for grasping the underlying logic of homeostatic regulation. We take as a starting point the formalization provided by control theory, which allows modeling the dynamics of regulatory mechanisms without explicitly considering their intricate mechanistic details. We adapt this formalization to address singular features of homeostatic systems such as the particular nature of biological signals.

Within the control theory framework, control signals typically express the distance between the current value of the regulated variable and its set point. This reference value is normally predefined and used as an input for the control loops that govern the system’s behavior [16, 21–23]. In contrast, biological control signals often take the form of molecules that must be continuously synthesized and degraded, and operate within complex molecular networks. This complexity conditions the conveyed information, and opens new opportunities for modulating their homeostatic effects.

Our modeling framework naturally integrates the functional and regulatory aspects of homeostatic systems, incorporating the particularities of biological signals. From this approach, it is possible to formulate simple abstract schemes that capture the regulatory logic of well-known homeostatic mechanisms. Despite their simplicity, these basic models provide valuable insights into key aspects of homeostatic systems. Specifically, they suggest that the notion of a predefined set point may not be essential for modeling biological regulation, and could potentially lead to misinterpretations about the role and the fundamental principles underlying homeostasis, one of the fundamental processes that sustain life.

## Results

### External and homeostatic flows

In this section, we formulate a simple modeling framework to simulate the dynamics of homeostatic control mechanisms. To do that, we use the graphic formalism of systems dynamics, in which variables are represented as boxes, and flows that affect the system’s variables are shown as arrows (see Figs. 1.A,B) [24]. A detailed introduction to the application of this formalism to biological homeostasis can be found in reference [14].

The model shown in Fig. 2.A can be interpreted as describing the behavior of a given molecule *M* whose concentration changes under the influence of an inflow and an outflow. The dynamics of this simple system depend on the particular form taken by these flows. For simplicity, let us consider the following equation:

**Figure 2.**
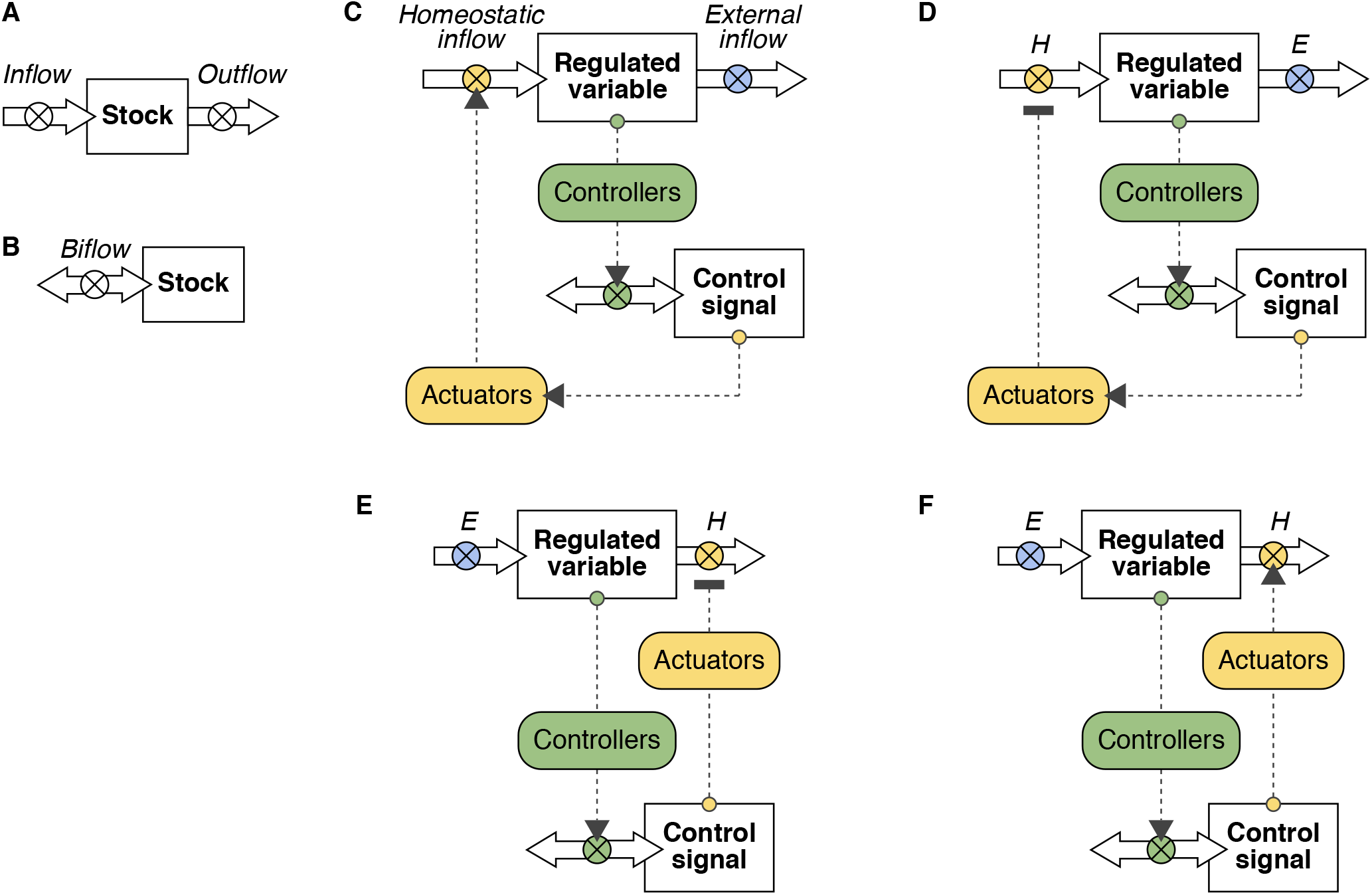
Stock and flow representation of simple homeostatic systems. A, B) Basic elements of system’s dynamics models. Stocks represent variables and solid arrows indicate inflows (positive flows) and outflows (negative flows), processes that increase and decrease the value of variables respectively (A). Biflows allow for either positive or negative flows from a stock. The direction of the flow is determined by the biflow’s sign (B). C,D) Demand-driven homeostatic systems. The value of the regulated variable changes under the influence of an unregulated external outflow. Control signals may balance this effect by upregulating (C) or inhibiting (D) a homeostatic inflow. In turn, the value of the regulated variable is used by controllers to increase or decrease the levels of the control signal. E,F) Supply-driven homeostatic systems. In these models, the value of the regulated variable is determined by the balance between an external inflow and a homeostatic flow, inhibited (E) or upregulated (F) by the control signal. Controllers and actuators are implicitly considered through their resultant influence on the dynamics of the system. Dashed arrows represent the flow of information. Blunt and pointed arrows indicate inhibition and upregulation respectively. H: homeostatic flow; E: external flow.

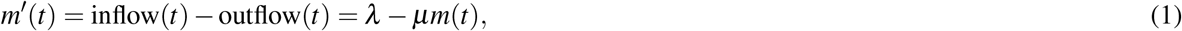

where *m*(*t*) is the concentration of *M* at time *t*, and *λ* and *μ* are positive parameters. In this model, *M* is produced at a constant rate *λ* and disappears exponentially at a constant rate *μ*. Although this system is not regulated, it reaches a stable steady state given by

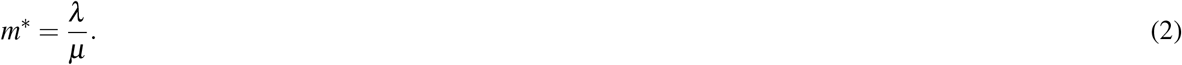

This basic example shows that regulation is not necessary for the system’s stabilization. However, the absence of regulation means that the concentration of *M* at the steady state (denoted by *m*^*\**^) can fluctuate outside the system’s control, potentially accumulating indefinitely or vanishing as the rates *λ* and *μ* approach zero.

Regulation can be implemented in model 1 by means of the following equations:

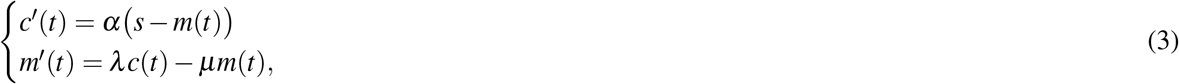

where *m*(*t*) and *c*(*t*) are the concentration of molecule *M* and a control signal *C* at time *t* respectively, and *α, s, λ*, and *μ* are positive parameters.

In the terminology of engineering control theory, parameter *s* denotes the system’s predefined set point, and the control signal *C* expresses the system’s error, i.e. the difference between the set point and the current value of the regulated variable (*m*(*t*)). Specifically, the concentration of the control signal increases when this difference is positive and decreases otherwise. In turn, the control signal acts by promoting the synthesis of molecule *M* (term *λc*(*t*)), which is consumed at a constant rate *μ* as in equations 1.

The main consequence of this regulatory mechanism is that the steady-state concentration of *M* is no longer affected by changes in consumption (parameter *μ*):

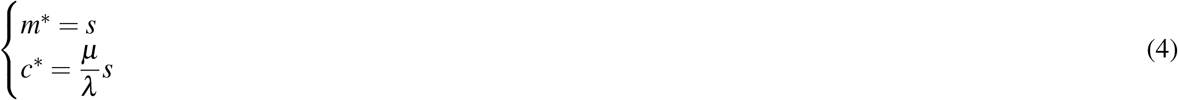

In the previous model, the control signal upregulates the production of *M*. Alternatively, model 1 can be regulated by inhibiting this production:

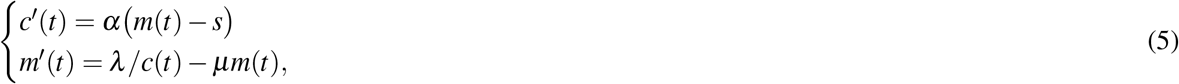

In this case, the synthesis of molecule *M* is greater for lower levels of the control signal. Reciprocally, these levels increase when the value of the regulated variable exceeds the set point (*s*) and decrease otherwise. The steady state of this regulated system is now given by

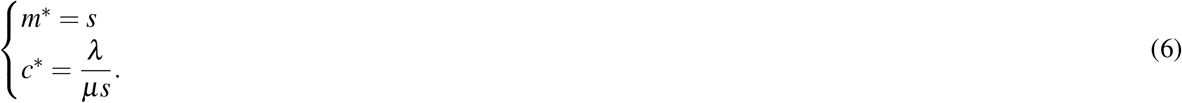

As with model 3, regulation eliminates the influence of the rate of consumption of the steady-state value of the regulated variable.

The logic of the previous regulatory mechanisms is represented in Figs. 2.B and C. In these systems, only the production of *M* is regulated. It is important to stress that this does not imply that consumption is unregulated, only that its regulation does not depend on the control signal *C*. We will label as homeostatic the flows that are regulated by the control signal (in these cases, the inflow), and as external those that are not regulated (in these cases, the outflow).

The external and homeostatic nature of inflows and outflows can be reversed in other control mechanisms. Let us consider, for instance, the following equations:

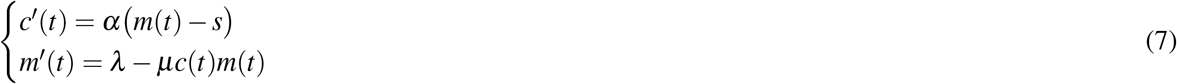

In this case, the control signal regulates the removal of the regulated molecule *M* (term −*μc*(*t*)*m*(*t*)), which is produced at a constant rate *λ*. To do that, the levels of the control signal (*c*) increase if the concentration of *M* is above the set point (*σ*) and decrease otherwise. In this model, homeostatic and external flows correspond to the outflow and the inflow respectively (see Fig. 2.C).

The regulation of a system with an external inflow can also rely on the inhibition of the homeostatic outflow. Such a regulatory mechanism can be modeled, for instance, as:

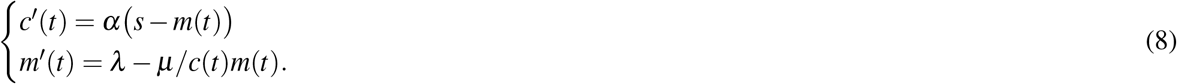

The steady state of these systems are, respectively:

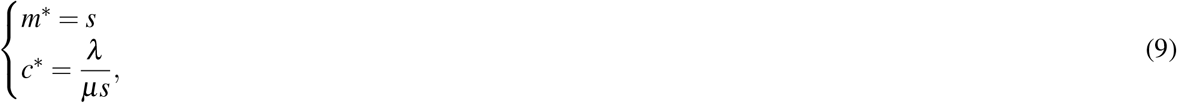

and

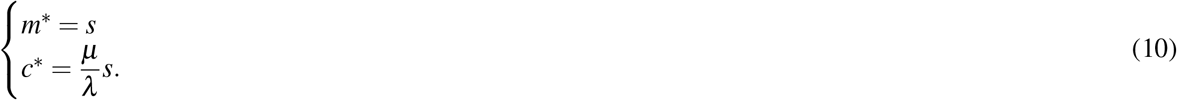

Again, as a consequence of regulation, the steady-state concentration of *M* in these systems is not affected by the magnitude of the external inflow (i.e. by parameter *λ*).

Following the terminology presented in [14], we will label models 3 and 5 as demand-driven systems, and models 7 and 8 as supply-driven systems. In the following section, we show that this approach provides a natural framework to model biological homeostatic systems.

### Demand- and supply-driven homeostatic systems

The models introduced in the previous section capture the regulatory logic of well known homeostatic mechanisms. For instance, the regulation of intracellular and systemic iron levels can be described as demand-driven homeostatic systems. Aconitase regulates intracellular oxygen homeostasis according to the following mechanism. When cellular iron is abundant, aconitase incorporates iron-sulfur clusters, becoming active as an enzyme that catalyzes the conversion of citrate into isocitrate [25]. Conversely, under conditions of iron deficiency, aconitase loses its iron-sulfur clusters and undergoes a conformational change that renders it inactive as an enzyme [26]. In this inactive form, aconitase transforms into an iron regulatory protein (IRP) that binds to mRNA transcripts of various genes involved in iron homeostasis, increasing cellular iron uptake and storage [26].

Within the modeling framework shown in Fig. 2, the IRP acts as a control signal that upregulates the homeostatic inflow of iron into the cell to compensate for the iron consumption by intracellular proteins and enzymes. Reciprocally, cellular iron promotes the disappearance of IRP by facilitating its conversion into aconitase, thus reducing its activity as a post-transcriptional regulator (Fig. 3.A).

**Figure 3.**
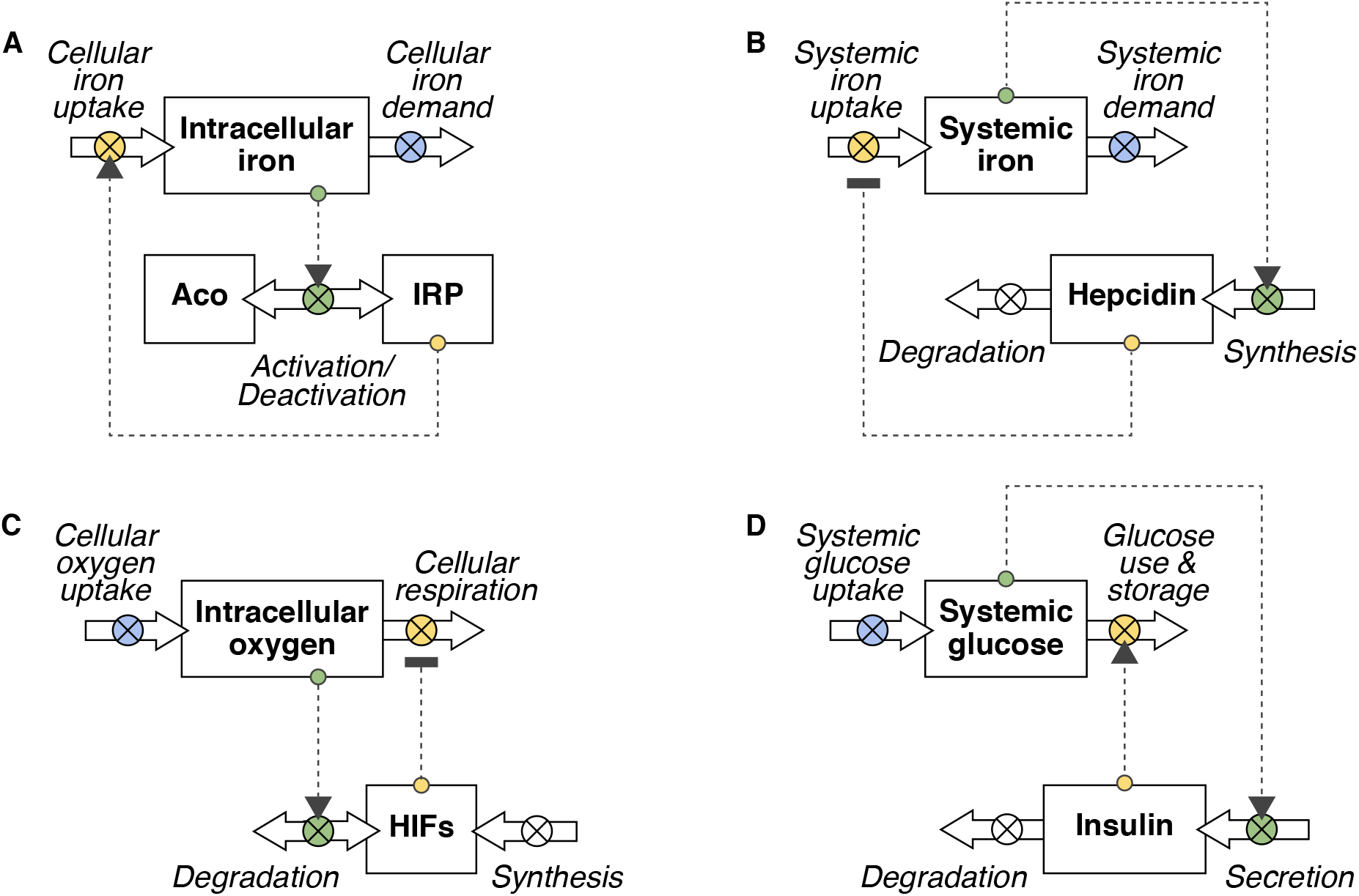
Examples of supply- and demand-driven homeostatic systems. A,B) Demand-driven homeostatic mechanisms equivalent to the abstract models shown in Figs. 2.C,D. C,D) Supply-driven homeostatic mechanisms analogous to the models in Figs. 2.E,F. For simplicity, actuators and controllers are not shown. Flows regulated by controllers are shown in green, and those governed by actuators in yellow. External flows are shown in blue. Dashed arrows represent the flow of information. Blunt and pointed arrows indicate inhibition and upregulation respectively.

Systemic iron homeostasis, on the other hand, is regulated by hepcidin. This hormone controls systemic iron absorption from the gut by regulating the activity of ferroportin, an iron exporter located on the surface of intestinal epithelial cells [27]. To that end, hepcidin binds to ferroportin and induces its internalization and degradation, thereby reducing iron absorption from the gut [28]. Hepcidin also reduces iron release from macrophages, which play a crucial role in recycling iron from senescent red blood cells [29].

Hepcidin is primarily synthesized and secreted by the liver in response to systemic iron availability [30]. High iron levels stimulate hepcidin production, leading to decreased iron absorption and increased sequestration within cells [30]. Conversely, low iron levels suppress hepcidin production, allowing for increased iron absorption and release into circulation [30]. In this example, hepcidin is the control signal that regulates systemic iron availability. To that end, it controls the homeostatic inflow of iron into the circulation and its synthesis scales proportionately with iron levels (see Fig. 3.B).

The regulation of intracellular oxygen by Hypoxia-inducible factors (HIFs), and that of systemic glucose by insulin are examples of supply-driven homeostatic systems. HIFs are dimeric transcription factors composed of two subunits, HIF-*α* and HIF-*β*, that regulate the expression of multiple genes involved in the cellular use of oxygen [31]. In particular, HIFs target-genes inhibit aerobic respiration and promote anaerobic pathways of energy production, thus reducing the cellular demand for oxygen [32]. Within the context of intracellular oxygen levels, oxygen uptake is an external flow, since it depends on factors (such as the local topology of the circulatory system, the number of red blood cells, or the heart and respiratory rates) that are not directly controlled by HIFs operating in an individual cell.

Reciprocally, oxygen facilitates the degradation of HIFs. Prolyl hydroxylase domain enzymes (PHDs) use oxygen to hydroxylate specific residues in the HIF-*α* subunit, thus marking it for proteasomal degradation [33]. This prevents the dimerization of HIF-*α* and HIF-*β* in the nucleus and the subsequent expression of HIFs target genes [33]. Consequently, HIFs can be regarded as a control signal governing the homeostatic outflow of oxygen in response to cellular oxygen uptake, with oxygen levels directly promoting their degradation (Fig. 3.C).

Finally, insulin promotes the removal of glucose from the blood, preventing an excess of glucose following dietary intake [34]. Insulin is produced by pancreatic *β* -cells in response to high glucose levels, and is removed from the circulation by the liver [35]. Therefore, insulin is a control signal that regulates the homeostatic outflow of glucose from the blood. In turn, its synthesis and secretion into the bloodstream increases with blood glucose levels (Fig. 3.D).

In this section, we have seen that the modeling approach introduced in the previous section provides a unifying framework to study homeostasis. From this perspective, homeostatic systems that are apparently very different can be understood as minor variations of common underlying principles (Fig. 3). We will next show that this functional aspect of homeostasis can provide valuable insights into key aspects of biological regulation.

### Set points and settling points

The goal of homeostatic regulation is widely understood as maintaining the value of key physiological variables within an adequate range [5, 17]. Changes in these variables are usually considered as perturbations that move the system away from its optimal steady state [5, 36]. Accordingly, homeostatic control systems are often described as stress response mechanisms that bring the perturbed system back to normality [15–18, 37].

However, biological variables may continuously change as a result of normal physiological activities. For instance, fluctuations in the intracellular levels of glucose or oxygen during routine cellular metabolism are the unavoidable consequence of healthy cell function and not necessarily a source of physiological stress. In fact, a major role of homeostatic regulation is to ensure that ordinary activities can proceed normally, preventing both the excess and the deficit of key molecules, which could compromise the function of a given biological system.

This ‘routine’ aspect of homeostasis is explicit in the models presented in Figs. 2.B-E and in Fig. 3. The view of regulation as driven by the supply or the demand of the regulated variable naturally accounts for the dynamics of the system, as shown in models 3 to 8, and provides a functional interpretation for the homeostatic role of control signals. This role would consist in adjusting the production of regulated molecules to compensate for an external demand (as in models 3 and 5), or modulating their removal from the system to balance an external input (as in models 7 and 8). As discussed earlier, regulation minimizes the effect of external flows on the steady-state of these models, contrasting with the unregulated scenario (see equations 2). This is achieved by explicitly incorporating the set point (*s*) into the systems’ dynamics as a parameter. The necessity of a predefined reference value to drive biological regulation remains a topic of debate [14]. In fact, homeostatic variables may reach a steady state even in the absence of a predefined reference value [38]. In such scenarios, the steady-state value of the regulated variable emerges as an output of its dynamics rather than being explicitly included as an input parameter in the system (see e.g., model 1). These emergent steady states have been termed settling points, in contraposition to set points [14, 38, 39].

It has also been argued that, while the set point serves as a useful abstract concept for describing homeostatic regulation, it may not fully capture the underlying biological mechanisms [8]. In this regard, the dynamics of the control signal in models 3 to 8 are directly influenced by the difference between the current value of the regulated variable and the predefined reference value *s*. The levels of the control signal increase or decrease based on whether the regulated variable is below or above the set point. This mode of action would require a mechanism capable of regulating both the production and removal of the control signal in response to changes in the regulated variable.

Such regulatory strategy is apparent in intracellular iron homeostasis, where iron availability simultaneously determines the activation and deactivation of the control signal (Fig. 3.A). However, in other biological scenarios, the synthesis and degradation of the control signal may not rely on the same mechanisms and may not both depend on the value of the regulated variable. This variability is illustrated in Figs. 3.B-D, where the regulated variable controls either the synthesis/secretion of the control signal (Figs. 3.B,D) or its degradation (Fig. 3.C), but not both simultaneously. For example, systemic iron levels only regulate the synthesis of hepcidin in the liver and its subsequent secretion into the circulation [40], whereas intracellular oxygen participates in the degradation of HIF-*α* but not in its synthesis [41].

Hence, from a mechanistic standpoint, the behavior of control signals in these homeostatic systems may not respond to the dynamics outlined in models 3 to 8. These examples underscore a fundamental aspect of biological control signals. Often, they are molecules subjected to a continuous turnover of synthesis and degradation, a process that may not conform to the logic of a predefined set point. Therefore, understanding the function of biological regulation requires the explicit consideration of control signal dynamics, a point that may not be obvious in control theory approaches to model homeostatic systems (see Fig. 1).

In the following sections, we will argue that the dynamic interplay between control signals and regulated variables is crucial to explain the nature of homeostatic steady states and their role in biological regulation. To that end, we will focus on the model illustrated in Figs. 3.C,D.

### The central role of control signals’ dynamics in biological homeostasis

Fig. 3.D represents the basic logic underlying blood glucose regulation by insulin. Insulin production and secretion increase with the levels of glucose in the blood. In turn, insulin promotes the removal of glucose from the circulation by increasing cellular glucose uptake and storage. The dynamics of this regulatory mechanism can be modeled as

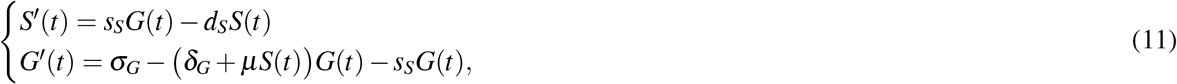

where *S*(*t*) and *G*(*t*) represent the concentration of insulin and glucose in the blood. Parameters *s*_*S*_ and *d*_*S*_ denote insulin synthesis and degradation rates respectively. The external glucose inflow is labeled as *σ*_*G*_ and the baseline rate of glucose consumption as *δ*_*G*_. The glucose homeostatic outflow controlled by insulin is modeled by the term *μS*(*t*)*G*(*t*) for a positive parameter *μ*.

It is important to note that blood glucose regulation does not exclusively rely on insulin. This is a highly complex process that involves many other signals and mechanisms operating both at systemic and cellular levels. It is patent that such complexity is beyond the reach of model 11. Indeed, this model is not intended as a detailed explanation of blood glucose homeostasis but as a simple tool to gain insight into key features of general homeostatic systems.

Blood glucose levels remain at relatively stable values (approximately 5 mM in human adults [4]), adequate for meeting cellular energy demands while preventing glucose excess. This homeostatic state is commonly attributed to the interplay between insulin and glucose shown in Fig. 1.B. However, this control mechanism (as formalized in Fig. 3.D) does not necessarily ensure a fixed steady state for blood glucose levels, since it cannot prevent the increase in blood glucose levels in scenarios where glucose uptake rises or glucose consumption diminishes, as shown in Figs. 4.A and 4.B, respectively. The rise in blood glucose levels under insulin regulation is notably less pronounced compared to the unregulated system (an effect that depends on the rates of insulin synthesis and degradation) but it remains unbounded (Fig. 4.A).

**Figure 4.**
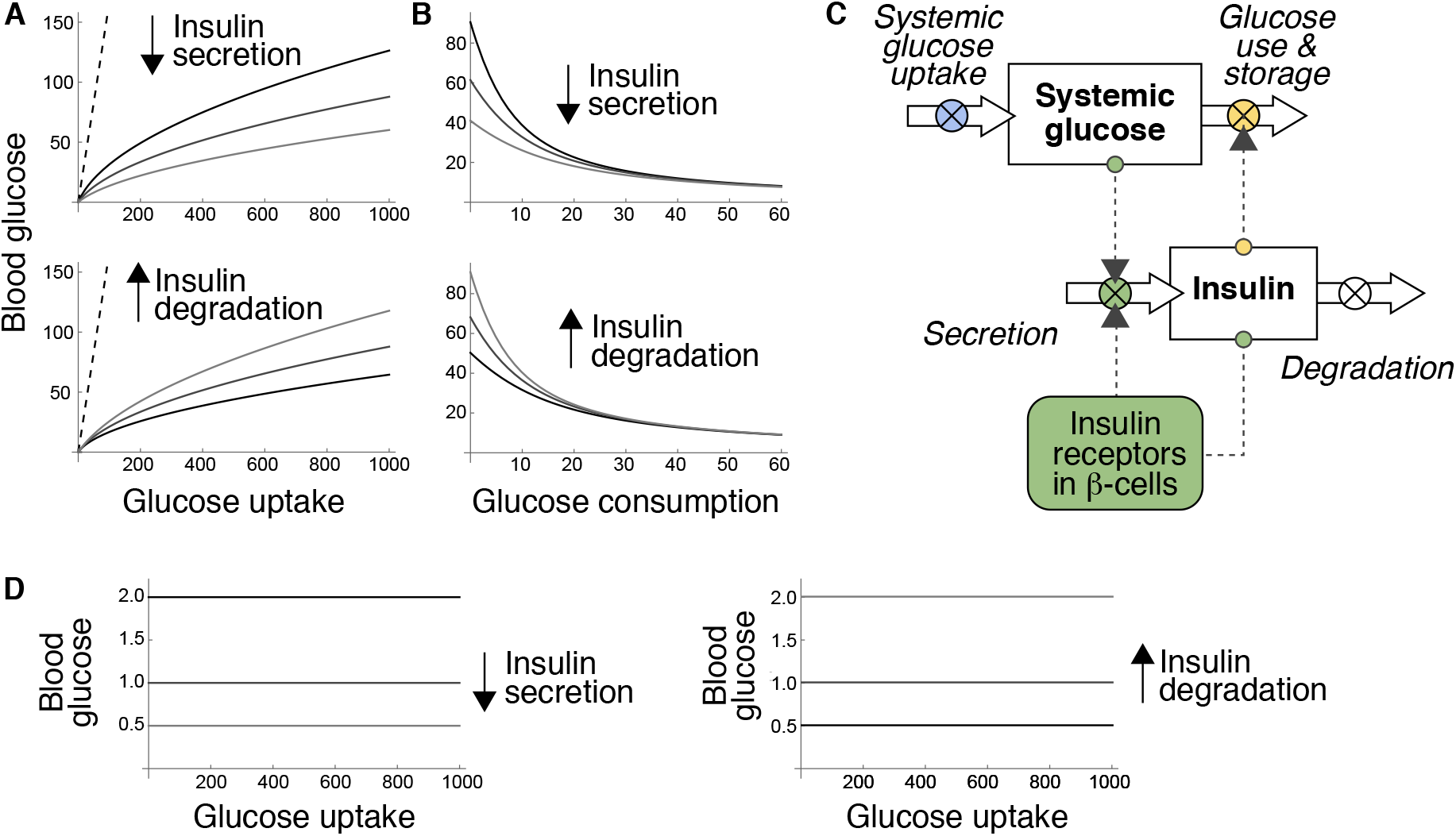
Models of blood glucose regulation by insulin. A) Steady-state value of blood glucose as a function of glucose uptake according to equations 11 for different rates of insulin secretion (upper) and degradation (lower). Dashed lines correspond to the unregulated system. B) Steady-state value of blood glucose as a function of glucose consumption according to model 11 for different rates of insulin secretion (upper) and degradation (lower). C) Diagram of a supply-driven model of insulin regulation including the paracrine effect of insulin on its own secretion. D)Steady-state value of blood glucose as a function of glucose uptake according to model 12 for different rates of insulin secretion (left) and degradation (right). Parameter values: *δ*_*G*_ = 0.6 and *μ* = 0.1. Scenarios of insulin secretion: *s*_*S*_ = 1, 2, 4 and *d*_*S*_ = 2. Scenarios of insulin degradation: *d*_*S*_ = 1, 2, 4 and *s*_*S*_ = 2. Dashed lines in A and B correspond to the unregulated model (*s*_*S*_ = 0, *d*_*S*_ = 0, and *μ* = 0). Parameters have been arbitrarily chosen to illustrate the dynamics of the models.

These results point to the need of additional mechanisms to ensure a fixed steady state for blood glucose levels. One such mechanism may involve the influence of insulin on its own production. Pancreatic *β* -cells, responsible for insulin production, express high levels of insulin receptors [42, 43], suggesting a possible autocrine regulation of insulin production [42, 44] (see Fig. 4.B). The potential consequences of insulin autocrine signaling remain controversial [45], and empirical evidence seems to support both inhibitory [46] and stimulatory [47] effects on insulin secretion. For the sake of the argument, we will assume that insulin exerts a positive effect on its own secretion (Fig-4.C).

Model 11 can be easily modified to include this autocrine effect. By way of example, let us consider the following equations:

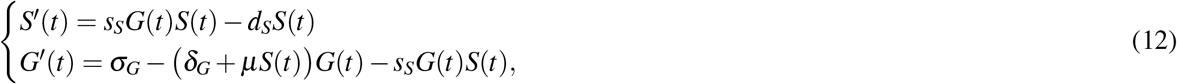

where insulin production is assumed to increase proportionally with insulin levels (term *s*_*S*_*G*(*t*)*S*(*t*)).

According to this model, the autocrine signaling of insulin on pancreatic *β* -cells would transform the system’s steady state, creating a fixed set point for blood glucose levels (Fig. 4.D). This set point would remain constant regardless of glucose uptake and demand, although it could be further modulated by changes in the rates of insulin synthesis and degradation (Fig. 4.D).

We insist that models 11 and 12 are not aimed at explaining blood glucose homeostasis in depth, but rather at elucidating fundamental aspects of biological regulation. Despite their simplicity, they offer valuable insights applicable to general homeostatic systems. Firstly, they suggest that describing control loops coupling the dynamics of regulated variables and control signals may not be sufficient to fully grasp the function of a homeostatic mechanism. For instance, the nature of the system’s steady state cannot be deduced from the diagrams in Figs. 1 and 4.C. This feature critically depends on the specific mechanisms governing the synthesis and degradation of the control signal and the dynamics they entail.

Secondly, equations 12 demonstrate that the same mechanisms of signal production and destruction may lead to either fixed or variable steady states, depending on the rates of control signal production and degradation (Fig. 4.D). Importantly, they also show that a regulated system can achieve a fixed steady state without relying on a predefined reference value. This behavior can be encoded in the dynamics of the control signal without the need for an explicit set point.

### Homeostatic regulation beyond set points

In this section, we will show that ensuring a fixed steady state is not always the goal of homeostatic systems. This is the case of intracellular oxygen regulation by HIFs. As discussed earlier, this regulatory mechanism is a supply-driven homeostatic system. Individual cells have little control over their oxygen supply, primarily determined by the tension of oxygen in their surrounding interstitial fluid [48]. To regulate intracellular oxygen levels, HIFs control the cellular consumption of oxygen, inhibiting aerobic respiration and facilitating anaerobic pathways of energy generation such as fermentation [49]. Reciprocally, oxygen levels determine the rate at which HIFs are degraded in the cell’s cytoplasm [41].

The dynamics of this regulatory scheme, corresponding to the system shown in Fig. 3.C, can be modeled as:

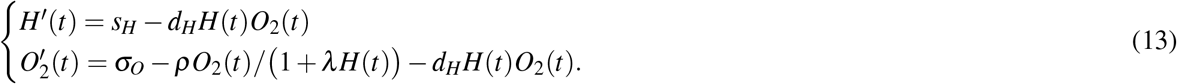

The abundance of oxygen and HIF-*α* in the cell at time *t* are denoted by *O*_2_(*t*) and HIF-*α*(t) respectively. This model assumes that HIFs transcriptional activity is proportional to HIF-*α* levels. Accordingly, the inhibition of respiration by HIFs is modeled as 1*/*(1 + *λ H*(*t*)), for a positive parameter *λ*. The rates of HIF-*α* synthesis and degradation are denoted by *s*_*H*_ and *d*_*H*_ respectively; *ρ* is the rate of oxygen consumption in respiration; and, finally, *σ*_*O*_ represents the external inflow of oxygen into the system, which is given by:

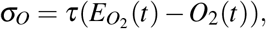

where 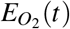 denotes the extracellular oxygen levels at time *t* and *τ* is a positive parameter.

This simple model predicts the empirically observed variation of HIFs expression with extracellular oxygen tensions: HIF-*α* levels are low in normoxia and increase exponentially as oxygen availability decreases [50, 51] (Fig. 5.A). According to equations 13, this pattern of expression implies that the steady state of intracellular oxygen is not fixed. Denoting by *H*^*\**^ and 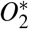 the steady states values of HIF-*α* and oxygen respectively, we have that 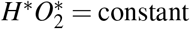.

**Figure 5.**
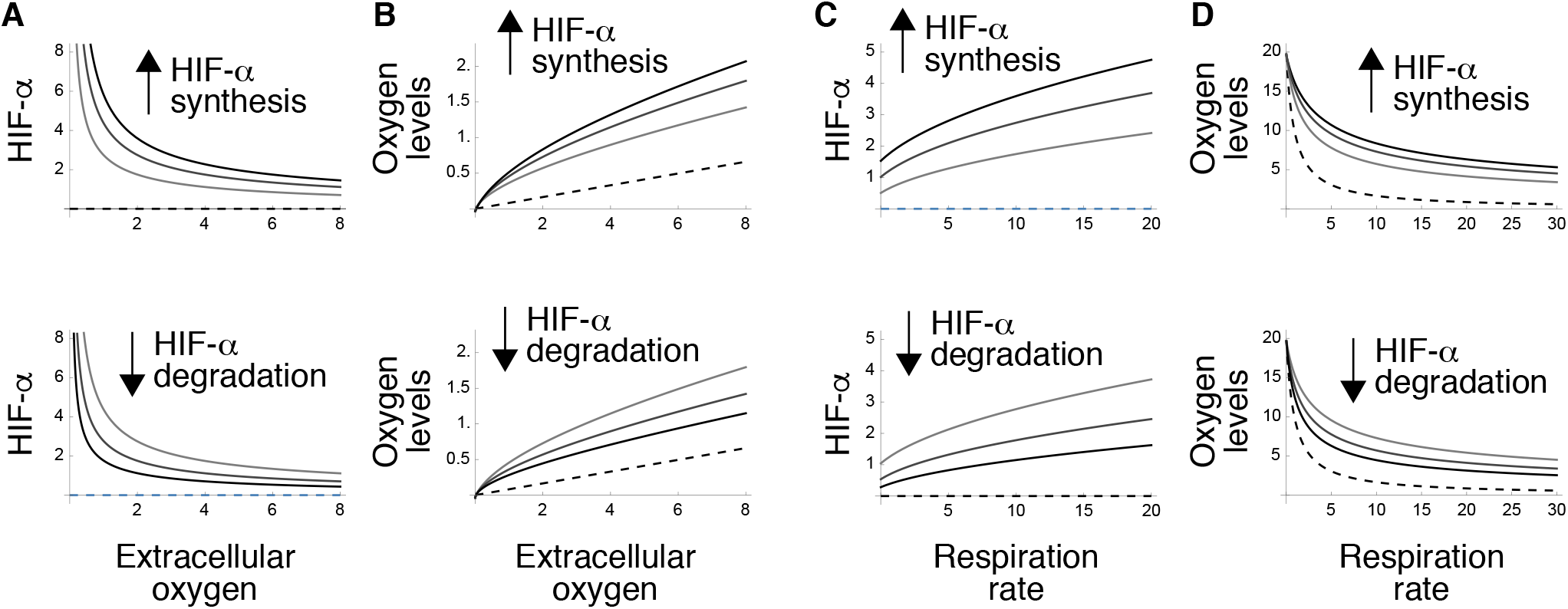
Regulation of intracellular oxygen homeostasis by Hypoxia-inducible factors (HIFs) according to model 13. A) Steady-state expression of HIF-*α* as a function of extracellular oxygen levels for different values of HIF-*α* synthesis (upper) and degradation (lower). B) Steady-state value of intracellular oxygen as a function of extracellular oxygen availability for different values of HIF-*α* synthesis (upper) and degradation (lower). C) Steady-state expression of HIF-*α* as a function of respiration rate for different values of HIF-*α* synthesis (upper) and degradation (lower). D) Steady-state value of intracellular oxygen as a function of respiration rate for different values of HIF-*α* synthesis (upper) and degradation (lower). Dashed lines correspond to the unregulated system. Parameter values: *ρ* = 10, *λ* = 2, and *τ* = 0.9. Scenarios of HIF-*α* synthesis: *s*_*H*_ = 0, 100, 200, 300 and *d*_*H*_ = 0.01. Scenarios of HIF-*α* degradation: *s*_*H*_ = 200, and *d*_*H*_ = 0.01, 0.02, 0.03. Parameters have been arbitrarily chosen to illustrate the dynamics of the models.

Therefore, steady-state intracellular oxygen levels are variable, increasing with extracellular oxygen availability (Fig. 5.B). HIFs activity is also affected by the cellular respiration rate, denoted as *ρ* in the previous equations (Fig. 5.C). As a consequence, HIFs regulation ensures higher intracellular oxygen levels as compared to the unregulated system in circumstances of increased metabolic demand (Fig. 5.D). This example clearly illustrates the homeostatic effects of the rates of control signal’s synthesis and degradation (Fig. 5).

HIFs also demonstrate that not all regulatory mechanisms are aimed at ensuring a fixed steady state for the system. Even if it is not fixed, the system’s steady state is not a passive result of the balance between oxygen uptake and respiration. It is actively determined by HIFs, which would not be merely sensors of intracellular oxygen [52], but regulators of a homeostatic outflow, functionally equivalent to insulin.

## Discussion

In this work, we introduce a modeling framework aimed at understanding the behavior of homeostatic systems. This framework integrates elements from control theory, adapting them to accommodate the unique features of biological regulation. Homeostatic mechanisms are particular instances of control systems and can thus be modeled using the analytical tools developed in engineering control theory. This approach provides a valuable description of homeostatic systems in terms of regulated variables, control signals, controllers, or actuators [14]. Such abstraction is useful for studying homeostasis as it allows modeling complex biological processes (such as the action of actuators and controllers) through their overall effect on the system’s dynamics, without detailing their specific modes of action.

Nevertheless, biological systems and engineering control systems diverge in crucial aspects, and overlooking their differences may lead to misinterpretations of the fundamental principles governing homeostatic regulation. The application of the set point concept in homeostasis is a paradigmatic example of this issue. In engineering systems, set points serve as predefined references to control the behavior of regulated variables [13]. As a result, control systems tend to stabilize at the desired values [13]. Prototypical homeostatic variables (e.g. blood glucose or body temperature) exhibit a similar behavior, maintaining relatively constant values regardless of variations in external conditions [1, 14]. However, from this observation alone, it cannot be inferred that the dynamics of homeostatic variables are actually driven by a predefined set point. For this reason, although set points can be useful to effectively reproduce the dynamics of homeostatic systems [53], they may not shed light on the mechanisms underlying biological regulation [19].

Conversely, not all regulated variables exhibit fixed or steady states [8, 19]. For instance, intracellular oxygen levels vary with extracellular oxygen availability despite HIFs regulation. Therefore, there exist regulatory mechanisms that cannot possibly rely on a predefined set point.

Fixed and variable steady states are often viewed as resulting from mutually incompatible processes: the former would be predefined and actively regulated by homeostatic mechanisms, while the latter would arise as passive results of the system’s dynamics [8, 17]. However, our models challenge this view, suggesting that this distinction is arbitrary. Both fixed and variable steady states may be encoded in the dynamics of control signals and emerge as outcomes of the system’s dynamics. The nature of the steady state would depend on the particular details of these dynamics.

Our approach also suggests that controlling the steady state of a system represents only a partial aspect of homeostasis. To gain a comprehensive understanding of biological regulation, it is essential to consider the function of regulated variables. From this standpoint, changes in the value of regulated variables should not be viewed as anomalous perturbations of the system but rather as a normal aspect of their physiological role. We argue that regulatory mechanisms can be naturally conceptualized as controlling either the supply or the demand of regulated variables [4]. This formalization highlights the functional analogies between a variety of homeostatic systems, ranging from the control of intracellular oxygen and iron to the regulation of systemic iron and glucose (Fig. 3).

Our interpretation of supply- and demand-driven systems differs from that provided in reference [4], where these control systems are presented as an alternative to homeostatic regulation. In [4], both supply- and demand-driven control systems are described as regulating the consumption of a given metabolite. The former would adjust consumption based on available supply, while the latter would modulate consumption in response to demand. In contrast, homeostatic control would entail the regulation of both supply and consumption to maintain the regulated variable close to its optimal state [4]. Additionally, the steady-state would be fixed in homeostatic systems, but it would vary in supply- and demand-driven systems as a consequence of regulation [4]. Under these definitions, the roles of supply/demand control and homeostatic regulation may complement each other, but they are essentially different [4].

While this modeling approach offers valuable insights [4], it does have limitations that constrain its explicative power. Firstly, we have seen earlier that regulation does not solely rely on adjusting consumption. For instance, the IRP or hepcidin regulate iron uptake at cellular and systemic scales respectively [26, 27], adjusting iron supply to meet the cell’s and the body’s demands. Furthermore, our model of insulin regulation suggests that simultaneous control of supply and demand is not necessary to ensure a fixed steady state for the regulated variable (see model 12). Our formalization of supply- and demand-driven control systems offers a broader perspective on biological regulation. According to this approach, the control of supply/demand of regulated variables and the nature of the system’s steady state represent two independent yet complementary dimensions of homeostatic regulation. The first dimension concerns the mechanisms underlying regulation itself. If a control signal is to govern the dynamics of a biological variable, then it must act upon the processes (supply and/or demand) that affect that variable. The second dimension is determined by the dynamics of the control signal, which dictate whether the system’s steady state is fixed or variable.

Supply- and demand-driven control are not alternative strategies to homeostasis. Instead, they represent two possible mechanisms of homeostatic regulation. On the other hand, homeostasis is not restricted to ensuring fixed steady states. This may be the main goal for some regulated variables, such as blood pH. Other regulatory systems, however, may prioritize the regulation of homeostatic flows. For instance, the role of HIFs is better understood as adjusting oxygen consumption based on its availability, regardless of the steady state of intracellular oxygen levels. Control signals can simultaneously regulate both aspects of homeostasis as shown by insulin, which maintains adequate blood glucose levels while ensuring metabolic glucose availability.

This work provides a unifying framework to study biological regulation. This approach can be easily extended to model more intricate regulatory systems, as shown in Supplementary Figs. S1 and S2. Furthermore, it identifies abstract, general principles underlying homeostatic systems that may greatly vary in their molecular details. These principles set the basis for a rigorous formalization of biological control and contribute significantly to advancing our understanding of homeostasis, one of the defining features of living systems.

## Acknowledgments

Cr.F.A. was partially supported by the MINECO grant PID2022-138187OB-I00.

