## Supplementary Material for "A functional approach to homeostatic regulation"

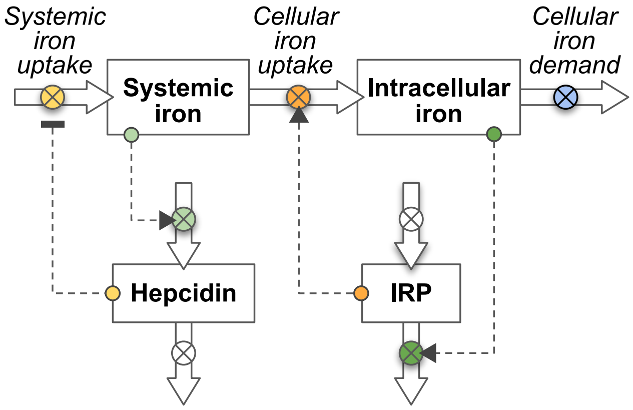


**Figure S1. Interaction between cellular and systemic iron homeostasis**. Intracellular iron homeostasis is a demand-driven mechanism that regulates the homeostatic uptake of iron from the blood. The iron consumed by the cells throughout the body constitutes the external demand that drives systemic iron homeostasis. Consequently, iron uptake is simultaneously a homeostatic flow from the viewpoint of cellular homeostasis, and an external flow in the context of hepcidin-regulated homeostasis. This example illustrates how the modeling framework presented in this work can be further developed to analyze homeostatic systems of increasing complexity.

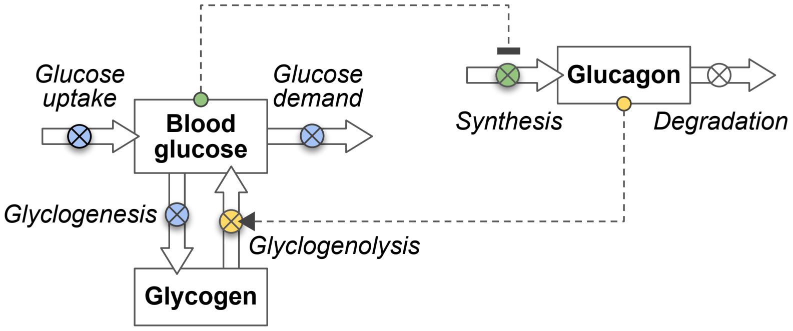


**Figure S2. Regulation of blood glucose homeostasis by glucagon.** Glucagon is produced by the $\alpha$-cells of the pancreas, and is removed from the circulation and metabolized in the liver and kidneys. This hormone promotes the breakdown of glycogen stored in the liver into glucose, a process known as glycogenolysis. Additionally, it stimulates gluconeogenesis, the synthesis of glucose from non-carbohydrate precursors such as amino acids and glycerol. These actions ensure that glucose availability in the bloodstream fulfills the body's energy needs, especially during fasting or periods of increased energy demand. Glucagon acts as a control signal that regulates the homeostatic inflow of glucose into the blood to counterbalance for glucose consumption in the body tissues. Conversely, high glucose levels in the bloodstream inhibit glucose secretion, a controller dynamics that differs from those outlined in main Fig. 3. The case of glucagon shows that the dynamics of the regulated variable may depend on several external flows and on additional variables such as glycogen. This dependence introduces a new layer of complexity that can be easily considered within the framework presented in this work.
